# NS2B-D55E and NS2B-E65D Variations are Responsible for Differences in NS2B-NS3 Protease Activities Between Japanese Encephalitis Virus Genotype I and III in Fluorogenic Peptide Model

**DOI:** 10.1101/2023.12.08.570834

**Authors:** Abdul Wahaab, Yan Zhang, Jason L. Rasgon, Lei Kang, Muddassar Hameed, Chenxi Li, Muhammad Naveed Anwar, Yanbing Zhang, Anam Shoaib, Ke Liu, Beibei Lee, Jianchao Wei, Yafeng Qiu, Zhiyong Ma

## Abstract

Japanese Encephalitis Virus (JEV) NS2B-NS3 is a protein complex composed of NS3 proteases and a NS2B cofactor. The N-terminal protease domain (180 residues) of NS3 (NS3(pro)) interacts directly with a central 40-amino acid hydrophilic domain of NS2B (NS2B(H)) to form an active serine protease. In this study, the recombinant NS2B(H)-NS3(pro) proteases were prepared in *E. coli* and used to compare the enzymatic activity between genotype I (GI) and III (GIII) NS2B-NS3 proteases. The GI NS2B(H)-NS3(pro) was able to cleave the sites at internal C, NS2A/NS2B, NS2B/NS3 and NS3/NS4A junctions that were identical to the sites proteolytically processed by GIII NS2B(H)-NS3(pro). Analysis of the enzymatic activity of recombinant NS2B(H)-NS3(pro) proteases using a model of fluorogenic peptide substrate revealed that the proteolytical processing activity of GIII NS2B(H)-NS3(pro) was significantly higher than that of GI NS2B(H)-NS3(pro). There were eight amino acid variations between GI and GIII NS2B(H)-NS3(pro), which may be responsible for the difference in enzymatic activities between GI and GIII proteases. Therefore, recombinant mutants were generated by exchanging NS2B(H) and NS3(pro) domains between GI and GIII NS2B(H)-NS3(pro) and subjected to protease activity analysis. Substitution of NS2B(H) significantly altered the protease activities, as compared to the parental NS2B(H)-NS3(pro), suggesting that NS2B(H) played an essential role in regulation of NS3(pro) protease activity. To further identify the amino acids responsible for the difference in protease activities, multiple substitution mutants including the individual and combined mutations at the variant residue 55 and 65 of NS2B(H) were generated and subjected to protease activity analysis. Replacement of NS2B-55 and NS2B-65 of GI to GIII significantly increased the enzymatic activity of GI NS2B(H)-NS3(pro) protease, whereas mutation of NS2B-55 and NS2B-65 of GIII to GI remarkably reduced the enzymatic activity of GIII NS2B(H)-NS3(pro) protease. Overall, these data demonstrated that NS2B-55 and NS2B-65 variations in hydrophilic domain of NS2B co-contributed to the difference in NS2B(H)-NS3(pro) protease activities between GI and GIII. These observations gain an insight into the role of NS2B in regulation of NS3 protease activities, which is useful for understanding the replication of JEV GI and GIII viruses.

## Introduction

Zika Virus (ZIKV), Dengue Virus (DENV) and Japanese Encephalitis virus (JEV) are mosquito borne members of the genus flavivirus and are resurging and emerging pathogens responsible for enormous disease burden. The flavivirus genome is ∼11kb single stranded, positive sense RNA comprising a large open reading frame encoding single protein precursor which is cleaved by a complex combination of cellular and viral encoded proteases during and immediately after translation. This cleavage generates three structural [Capsid (C), precursor membrane (prM), Envelope (E)] and seven nonstructural (NS1, NS2, NS2b, NS3, NS4a, NS4b and NS5) proteins necessary for viral genome replication [1–3]. JEV was first isolated in 1935 and since then has been classified to five genotypes (GI – GV) with geographic distribution in all continents except Antarctica, encompassing 67,900 human cases annually with a case fatality rate approaching 20-30%, and where 30-50% of survivors suffering from neurological sequelae [4–7].

JEV GI emerged in the 1990s and has progressively replaced GIII as the most frequently isolated JEV genotype from stillborn piglets, *Culex tritaeniorhynchus* mosquitoes, and clinical JE patients in Korea, China, Vietnam, India, Thailand, Taiwan and Japan [8–14]. It is suggested that JEV GI possibly competes with JEV GIII in same pig-mosquito cycle and has a transmission advantage over areas formerly dominated by GIII [11, 15]. Emerging JEV GI replicates more efficiently in wild birds, ducklings and cell cultures derived from *Aedes albopictus* mosquitoes compared to GIII [16–20] and shows similar infectivity in *Culex pipiens* [21]; however, it is unclear if replication efficiency of JEV GI vs JEV GIII occurs in *Culex tritaeniorhynchus* mosquitoes or/and pigs, which have crucial role in local JEV transmission. Additionally, experimental evidence linked with variance in viral genomic factors does not fully explain the occurrence of JEV GIII replacement during past two decades[11, 17, 22].

JEV encodes a protease complex comprising the viral NS3 protein and a NS2B cofactor which are requisite for generating functional viral particles. NS2B is an integral membrane protein comprising two hydrophobic and a central hydrophilic domain and plays a crucial role in viral replication [23, 24]. The central hydrophilic region of NS2B(H) comprising around 50 residues is essential for activation of NS3pro and accurate processing of viral polyprotein and is the principal determinant in substrate selection [25–27]. NS3 usually possess RNA Helicase, serine proteases, RNA triphosphatase and Nucleoside triphosphatase enzymatic activities [28].These two component encoded NS2B-NS3 proteases are involved in polypeptide processing, RNA replication, infectious particle assembly via enzymatic-independent or -dependent processes and thus it substitutions may augment viral replications in different hosts [29–31]. Moreover, this two-component protein harbors diverse strategies to escape host innate immunity [22, 32]. It has potential to cleave interferon stimulators and surface receptors tyrosine kinase. These interferon antagonistic abilities facilitate efficient viral replication, viral particle release, and neuroinvasion which contributes to augmented virulence and high mortality in laboratory animals [33–36].

In this study, we investigated the contribution of NS2B/NS3 proteases for their possible role in emergence of JEV GI over previously emerged GIII *in vitro*, using a fluorogenic peptide-based model. The methodology involved cloning, expression, and purification of active and inactive JEV GI and GIII Ns2B(H)-NS3(pro) serine proteases and obtaining numerical data on kinetic constants through fluorescence resonance energy transfer (FRET) modeling, using substrate sites suspected to be cleaved by viral serine proteases. The involvement of mutations in NS2B(H) hydrophilic and NS3(pro) protease domains with different proteolytic processing activities of GI and GIII was also determined using recombinant proteins. The identified genetic determinants will be crucial for selection of genes to monitor GI virus evolution, replication, and activities in natural transmission cycles.

## Material and Methods

### Virus stock, Cells and Antibodies

Japanese encephalitis virus GI strain SH7 (MH753129) and GIII strain SH15 (MH753130), isolated from *Cx. tritaeniorhynchus* and *An. sinensis* (respectively) in 2016 were used in this study [37, 38]. All JEV strains were plaque-purified three times and amplified in BHK cells at 0.1 MOI as described previously [37]. The fifty percent tissue culture infective dose (TCID50) was determined, and fresh virus suspensions were used in all experiments. Baby hamster kidney cells (BHK-21) cell line was obtained from the American Type Culture Collection (ATCC) and cultured in Dulbecco’s modified Eagle’s medium (DMEM, Invitrogen, GIBCO, Carlsbad, CA, USA) containing 10% fetal bovine serum (FBS) (Gibco, Thermo Fisher Scientific, Waltham, MA, USA) and 100 μg/ml streptomycin and 100 IU/ml penicillin at 37℃. The commercial antibodies used in this study included a GFP (D5.1) XP Rabbit monoclonal antibody (Cell Signaling Technology, Danvers, MA, USA), GAPDH (glyceraldehyde-3-phosphate dehydrogenase) monoclonal antibody (Proteintech, Chicago, IL, USA), monoclonal His-tag antibody (GeneScript, Piscataway, NJ, USA) and a self-generated antibody specific to JEV NS3 [39].

### Sequence Alignment

Amino acid sequences of Japanese Encephalitis Virus genotypes including GI-SH7 strain (GenBank no MH753129.1) and GIII-SH15 strain (GenBank no MH753130.1) were assembled from the NCBI database (http://www.ncbi.nlm.nih.gov). Sequence of recombinant NS2B(H)-NS3(pro) proteases and the predicted/anticipated cleavage sites at internal capsid (internal C), Cap/prM, prM/E, E/NS1, NS1/NS2A, NS2A/NS2B,NS2B/NS3, internal NS3, NS3/NS4A, internal NS4A), NS4A/NS4B and NS4B/NS5 intersections were aligned using SnapGene (GSL Biotech LLC, San Diego, CA, USA) and MegAlign (DNASTAR Inc, Madison, WI, USA) software.

### Construction of PTrcHis-A NS2B(H)-NS3pro

Total RNAs from BHK cells infected with JEV GI-SH 7 and JEV GIII-SH15 strains were extracted using TRIzol reagent (Thermo Fisher Scientific, Waltham, MA, USA) according to the manufacturer’s protocol. Reverse transcription was performed to make cDNA using PrimeScript RT reagent kit with gDNA Eraser (TaKaRa, Kyoto, Japan). Sequences encoding the C-terminal portion (amino acid 45 to 131) of NS2B and N-terminal portion (amino acid 1 to 185) of NS3 were amplified and inserted into the multiple cloning site of protein expression plasmid pTrcHisA (Thermo Fisher Scientific, MA, USA) through restriction sites XhoI and EcoRI. Sequence encoding the 96-120 NS2B residues was removed from recombinant pTrcHisA plasmid by PCR based site directed mutagenesis [40] using Pfu ultra II fusion HS DNA polymerase (Agilent, Santa Clara, USA) to generate recombinant plasmids expressing N-terminal hexahistidine tag fused active NS2B(H)-NS3(pro) proteases [41, 42]. The active recombinant NS2B(H)-NS3(pro) proteases were inactivated [43] by replacing a serine at residue 135 with an alanine residue. Point mutations in NS2B(H) regions of proteases were attained through PCR based site directed mutagenesis [40]. The resulting constructs were sequenced in both direction through plasmid universal primers (PTrcHis Forward/Reverse) with an ABI Prism 3730 DNA sequencer (Applied Biosystem, USA) at Invitrogen (Guangzhou, PR China).

### Construction of Recombinant PET Duet1 Co-Expressing Artificial GFP Substrate and NS2B(H)-NS3(pro)

Sequences encoding JEV GI (SH 7) active and inactive NS2B(H)-NS3 pro proteases were generated as described previously and cloned into the MCS1 (multiple cloning site 1) of dual protein expression plasmid pETDuet-1 (Novagen, Beijing, China) through restriction sites EcoRI and NotI. For artificial GFP substrate, sequence encoding anticipated cleavage sites i.e. Internal C, C/prM, prM/E, E/NS1, NS1/NS2A, NS2A/NS2B, NS2B/NS3, internal NS3, NS3/NS4A, internal NS4A, NS4A/NS4B and NS4B/NS5 intersections were cloned from JEV GI SH7 (Figure 1B, 1C) strain with particularly designed oligo dT primer pairs (Shanghai Sunny Biotec, Shanghai, China) by reverse transcriptase polymerase chain reaction (RT-PCR) and inserted between N-terminal (amino acid 1 to 173) and C-Terminal (amino acid 174 to 239) of GFP by overlap extension PCR. The obtained sequence was cloned to MCS2 (multiple cloning site 2) of pETDuet-1 using[44]. Sequence of twenty alanine residues was inserted between N-Terminal and C terminal part of GFP to generate a control artificial GFP substrate.

**Figure 1:**
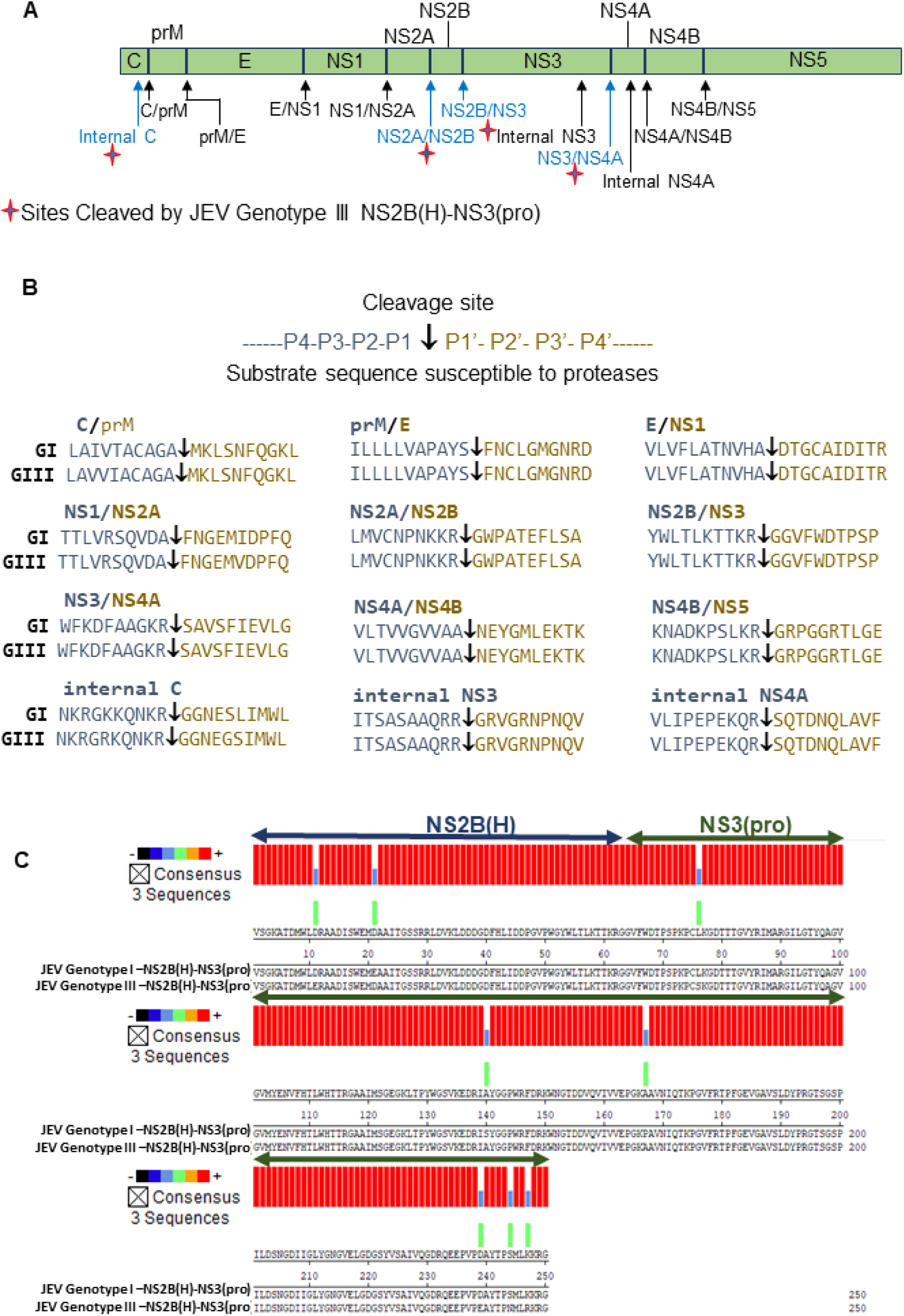
Japanese Encephalitis Virus two component NS2B(H)/NS3(pro) proteases and cleavage sites predicted to be proteolytically processed by them. **A)** Schematic diagram of JEV polyprotein with the predicted cleavage sites. Black arrowheads indicate the predicted cleavage sites and blue arrow heads represent the sites previously reported to be cleaved by JEV genotype III proteases (Wahaab et al 2021). (**B**) Sequence alignment of the predicted cleavage sites for JEV genotype I and III. Arrowheads indicates the predicted cleavage sites. **C)** Sequence alignment of two component JEV NS2B(H)/NS3(pro) proteases for JEV Genotype I (SH7) and Genotype III (SH15) showed eight amino acid substitutions between them which are shaded in blue. Homologous amino acids are shaded in red color.

### Detection of Cleavage Sites in competent *E. coli*

*E. coli* BL21 (DE-3) cells were transformed with recombinant pETDuet-1 expressing JEV GI NS2B(H)-NS3(pro) proteases and artificial GFP substrate and incubated at 37℃ until OD600 reached 0.6. Isopropyl b-D-1pthiogalactopyranoside (IPTG) was used to induce expression at a final concentration of 0.70 mM. The cells were harvested after 24 h incubation by centrifugation at 10000xg for 10 minutes at 4℃. The pellets were washed and resuspended in cold phosphate buffered saline (PBS 1X) and recentrifuged at 10000xg for 10 minutes at 4℃. The pellet was subjected to western blot analysis using specific antibodies as described previously [45]. The cleavage of artificial GFP substrate was detected by monoclonal anti-GFP antibody (GFP (D5.1) XP Rabbit). The expression of NS2B(H)-NS3(pro) was probed by antibodies specific to NS3 [39].

### Expression and Purification of NS2B(H)-NS3(pro) proteases

PTrc His-A vector harboring various JEV NS2B-NS3 proteases were propagated in *Escherichia coli* DH5 (TIANGEN Biotech, Beijing, China) and extracted using Plasmid Miniprep Kit (Corning, NY, USA). Extracted constructs were transformed to *E. coli* BL-21 (DE-3) cells for expression and cells were grown in 1000 ml LB medium comprising 100mg/ml ampicillin at 37℃ until the OD600 reached 0.6. The temperature was reduced to 18C and isopropyl b-D-1pthiogalactopyranoside (IPTG) was used to induce expression at a final concentration of 0.70mM. The cells were incubated for 14 hours at 18C. The cells were harvested by centrifugation at 8000xg for 10 minutes at 4℃. The pellets were washed and resuspended in cold phosphate buffered saline (PBS 1X) and recentrifuged at 8000xg for 10 minutes at 4℃. Pellet was kept on ice and resuspended in 40 ml Lysis buffer (0.1 M Tris-HCl, 0.3 M NaCl, PH 7.5, 10 mg ml-1 DNase, 0.25mg ml-1 lysozyme and 5mM MgCl2). Cells were kept at room temperature for 30 minutes and lysed on ice by sonication with an Ultrasonic Processor XL (Misonix Inc, NY, USA) Residues in solution were pelleted by centrifugation at 12000Xg for 30 minutes at 4℃ and soluble fraction was filtered through 0.22-micron filters (Merck Millipore Ltd, Cork, IRL). Supernatant was loaded to a 5 ml Nickel-Sepharose HisTrap Chelating column (GE Healthcare) pre-equilibrated with lysis buffer at flow rate of 1.0 ml per minute. The column was washed with five column volumes of buffer A (0.1M Tris HCl, pH 7.5, 0.3M Nacl, 20mM imidazole) and five column volumes of Buffer B (A (0.1M Tris HCl, pH 7.5, 0.3M Nacl, 50mM imidazole) respectively. Proteins were eluted with ten column volumes of elution buffer (0.1M Tris HCl, pH 7.5, 0.3M Nacl, 250mM imidazole) in fractions of 1.0 ml in DNA/protease-free Eppendorf tubes. Aliquots of 20ul from each tube were subjected to 15% SDS-page and Gels were stained with One-Step Blue, Protein Gel Stain, 1X (Biotium, Fremont, CA, USA) as per manufacturer instructions. Fractions containing NS2B/NS3 proteins were pooled and desalted through stepwise dialysis as described previously [42]. Purified NS2B-NS3 pro was concentrated to 1.0 mg ml-1 using centrifugal filter columns (Centricon 20 ml, 3000 MWCO, Millipore, USA). Protein concentration was determined with enhanced BCA protein assay kit (Beyotime, Shanghai, China) with bovine serum albumin as standard. Samples were stored in 50mM Tris-HCl, pH 9.0, glycerol (50% V/V) at 20C. Proteins were subjected to western analysis using Anti-Ns3 and His Tagged antibodies.

### Protease activity Assay

The assay of protease activity of purified NS2B(H)-NS3(pro) was carried out using previously reported fluorogenic peptides from JEV conserved cleavage sites NS2B/NS3 (Pyr-RTKR-amc) and NS2A/NS2B (Dabcyl-PNKKRGWP-(EDANS)G) synthesized by A+Peptide (Pudong, Shanghai, China) [42]. Assays were conducted on 96-well black, flat bottom, tissue culture treated polystyrene microplates (Corning Life Sciences, MA, USA) in total reaction volume of 0.1 ml containing 0.5 mM NS2B(H)-NS3(pro) proteases, assay buffer (50mM Tris HCl, pH9.5, 20% glycerol) and substrate. For kinetic assessments, fluorescence measurements were recorded at 20 minutes interval for 520 minutes through Multimode plate reader (Bio Tek) at an excitation wavelength(λ_ex_) of 360nm and emission wavelength(λ_em_) of 460 nm. Assay was carried out with three repeats for each group in two independent experiments at a constant temperature of 37℃. Inner filter effects of microplate reader were corrected as described in literature [46]. To determine the amount of AMC/fluorescence released, a standard curve was plotted with various value of kinetic data. No significant hydrolysis of peptide substrate was noted in dead NS2B(H)-NS3(pro) and/or groups without enzymes. Data was analyzed using GraphPad Prism version 7.00 for windows (GraphPad Software, La Jolla, California, USA)[47, 48].

### Homology Modeling

The image of NS2B/NS3 was created using SWISS-MODEL (www.swissmodel.expacy.org). The structure of NS2B/NS3 was downloaded from protein data bank and was assembled and analyzed with PyMOL (http://pymol.org) and SPDV (DeepView) software’s (https://spdv.vital-it.ch).

### Statistical Analysis

Statistical analysis was performed using GraphPad Prism version 7.0 (GraphPad Software, La Jolla, California, USA). Data were assembled as mean with standard deviation. Significant differences between groups were determined by Student’s t-test. A “p value” less than 0.05 (P<0.05) was considered as significant.

## Results

### Amino acid variation in cleavage sites and viral NS2B-NS3 proteases

JEV polyprotein is cleaved to generate functional proteins by a combination of host and viral proteases. The cleavage is anticipated to occur at connections between C/prM, prM/E, E/NS1, NS1/NS2A, NS2B/NS3, NS3/NS4A, NS4A/NS4B, NS4B/NS5 and sites of internal C, NS4A and NS3 (Fig. 1A). Alignment of prophesied cleavage site sequence revealed no variations among strains of GI and GIII (Fig. 1B). However, the two components viral NS2N(H)/NS3(pro) proteases involved in proteolytic processing of virus polyprotein exhibited eight mutations between GI and III (Fig. 1C). Two conserved mutations were spotted in hydrophilic domain of NS2B region i.e E55D and D65E with conservation rate of 90-100 %, and six mutations were spotted in protease domain of NS3 region i.e L14S, S78A, P105A, D177E, K185R with conservation rate of 90-100 % and S182 with a conservation rate of 58-89% (data obtained after alignment of 50 random represented strains from GI and GIII) [49]. Three mutations in NS2B-NS3 regions of JEV were previously reported to be involved in replication advantage of GI over GIII in avian cells at elevated temperatures [50]. However, mechanism behind that advantage still needs to be investigated making NS2B/NS3 a promising region to monitor virus evolution and genotype displacement.

### JEV GI and GIII NS2B(H)/NS3(pro) proteases exhibits identical cleavage patterns for proteolytic processing of JEV polyprotein

Studies have demonstrated that JEV protease activity mainly depends on association between NS3 and its co-factor NS2B, and that this two component NS2B-NS3 proteases expressed in *E.coli* are folded correctly with effective proteolytic activity [28, 42]. Therefore in our previous study we selected the prokaryotic cell model of *E. coli* to identify cleavage sites proteolytically processed by two component JEV GIII-SH15 proteases [51]. GI-SH7, which exhibits different 3D protein conformations (Fig.7B) and eight critical substitutions (Figure 1C) in the protease region compared to GIII-SH15 was engineered to a dual protein prokaryotic expression plasmid pETDuet-1 (Fig. 2A) to observe and compare the cleavage pattern of artificial GFP substrate containing cleavage site from GI-SH7 (Figure 2). Artificial GFP substrate, when expressed alone in *E. coli,* did not show any cleavage by *E. coli* proteases. When recombinant viral Ns2B(H)-NS3(pro) was co-expressed with all artificial substrates individually, a 21kDa band corresponding to N-terminal part of cleaved substrate was seen (Fig. 2B). The identified cleavage sites were validated by co-expression of all artificial substrates sites with inactivated/dead viral NS2B(H)-NS3(pro) which was generated by substitution of Serine at position 135 located in catalytic trait of NS3 with alanine (Fig. 2C). Cleavage of artificial GFP substrates was determined by western blot using antibodies specific to GFP and expression of viral proteases were determined by anti-NS3. Here our results demonstrated that GI-SH7 was able to cleave the sites at internal C, NS2A/NS2B, NS2B/NS3, and NS3/NS4A junctions and exhibited the same cleavage pattern as previously reported for SH-GIII [51], suggesting that viral protease substitution and protein conformation do not interfere with selection of cleavage sites for proteolytic processing and possibly play no role in genotype displacement.

**Figure 2.**
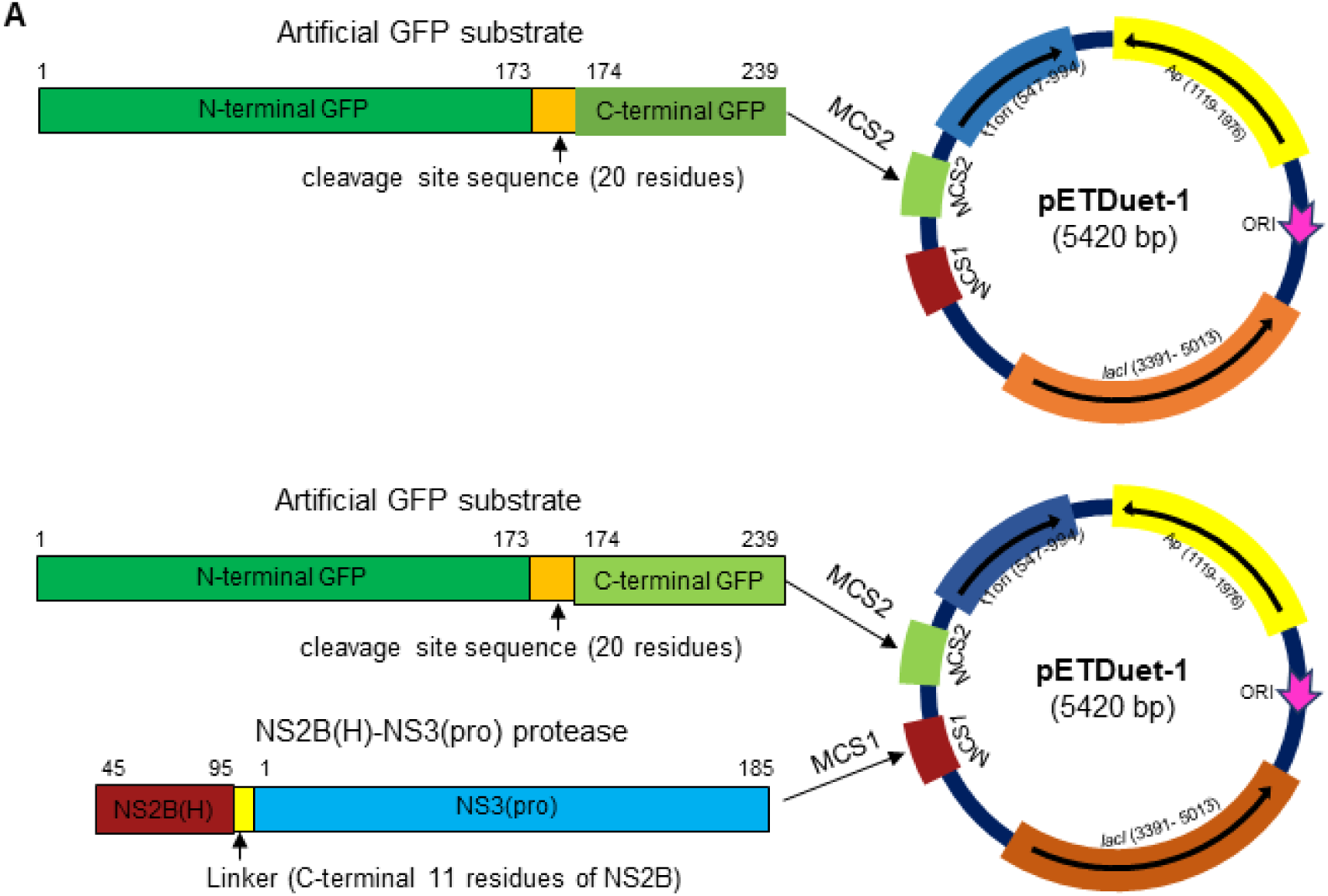
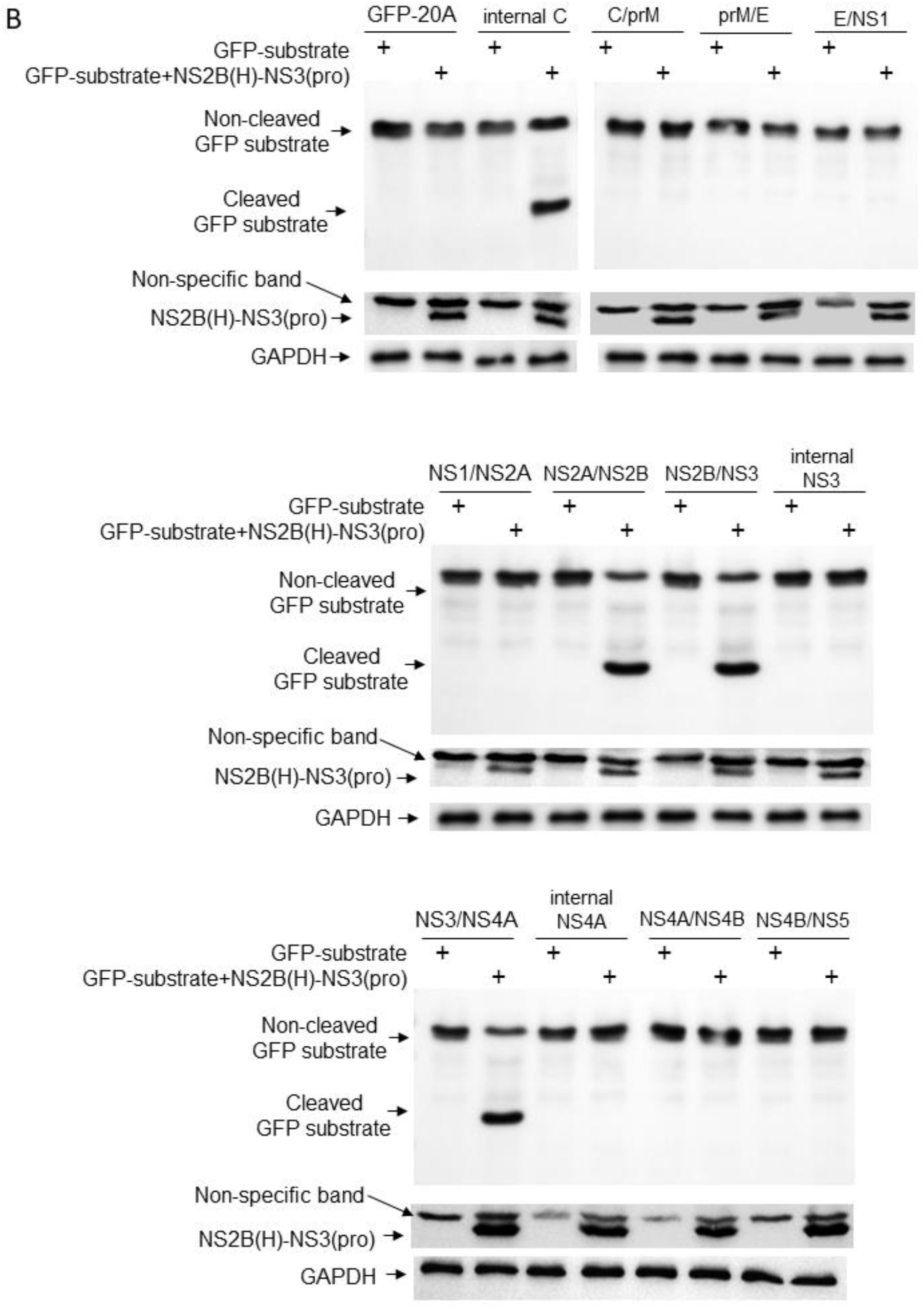
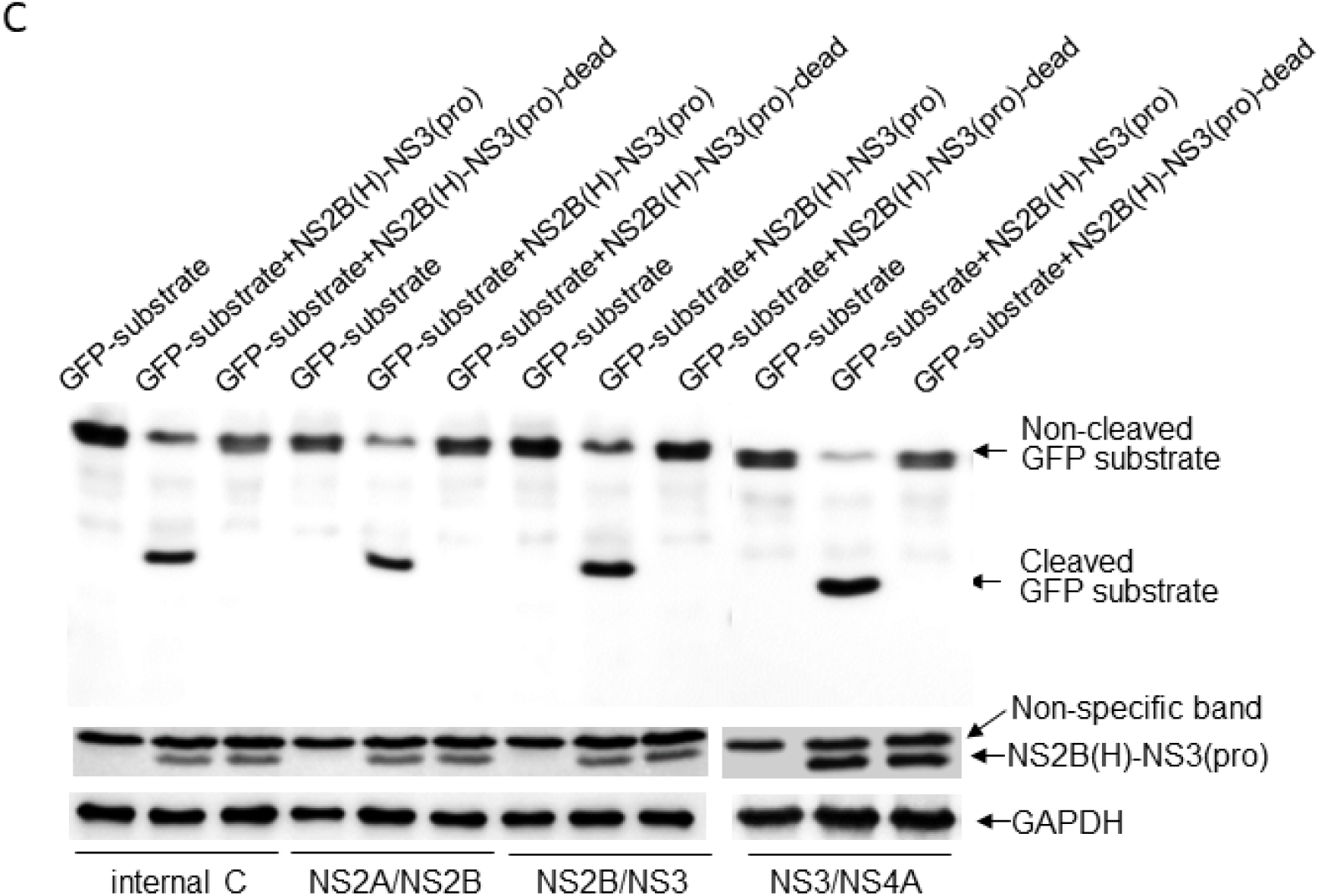
Detection of cleavage sites by JEV Genotype I NS2B(H)-NS3(pro) protease n *E. coli*. **A)** Schematic representation of recombinant plasmids **B)** *E. coli* cells were transformed with recombinant plasmid dually expressing and artificial GFP substrate or with recombinant plasmid expressing artificial GFP substrate alone. Cleavage of the artificial GFP substrate in E. coli was examined by western blot with antibodies specific to GFP. Expression of intact (non-cleaved). NS2B(H)-NS3(pro) protease was detected with antibodies specific to NS3. **C)** Identified cleavage sites were validated by transforming of E. coli cells with recombinant plasmid dually expressing active NS2B(H)-NS3(pro) protease and artificial GFP substrate (GFP-substrate+NS2B(H)-NS3(pro)), or inactive NS2B(H)-NS3(pro) protease and artificial GFP substrate (GFP-substrate+NS2B(H)-NS3(pro)-dead), or with recombinant plasmid expressing artificial GFP substrate alone. Cleavage of artificial GFP substrates in E. coli was examined by western blot with antibodies specific to GFP. Expression of intact (non-cleaved) NS2B(H)-NS3(pro) protease was detected with antibodies specific to NS3.

### Preparation of recombinant NS2B(H)-NS3(pro) proteases

Studies have demonstrated that JEV protease activity mainly depends on association between NS3 and its co-factor NS2B, and that this two component NS2B-NS3 protease expressed in *E. coli* is folded correctly with effective proteolytic activity [28, 42]. To analyze and compare the proteolytic processing activities of GI-SH7 and GIII-SH15 NS2B/NS3 proteases, the active and inactivated NS2B(H)-NS3(pro) proteases were structured and engineered into P-TrcHisA vector for expression in *E.coli* (Fig. 3A). SDS-PAGE and western immunoblotting of purified proteins was performed with antibodies specific to polyhistidine and JEV-NS3. SDS-PAGE revealed presence of three bands. The intact band of viral NS2B(H)-NS3(pro) proteases was seen at 36kDa. Whereas the autocleavage bands of NS3(pro) and NS2B(H) [42] were observed at 21kD and 10 kDa respectively for each active SH7 and SH15 proteases (Fig. 3B). However, only the intact NS2B(H)-NS3(pro) at 36Kda was seen for inactivated/dead NS2B/NS3 proteases of both strains (Fig. 3B), confirming its inactivity. The SDS-PAGE blots were quantified using imgaeJ software to determine self-cleavage percentage of NS2B-NS3 proteases of both strains. SH7 was self-cleaved with a percentage 89.9% of and SH15 with a percentage of 97.5%. The purified NS2B(H)-NS3(pro) proteases were further confirmed by western blot with antibodies specific to His-tag and NS3, repetitively. Distinct differences in intact anti-His and anti-NS3 expression was also seen between both strains (Fig. 3C). Overall, these observations suggest that GIII-SH15 shows high autocleavage rate of enzymatically active NS2B(H)-NS3(pro) proteases compared to GI-SH7. However, no autocleavage was seen for the dead/inactivated viral proteases of both strains.

**Figure 3:**
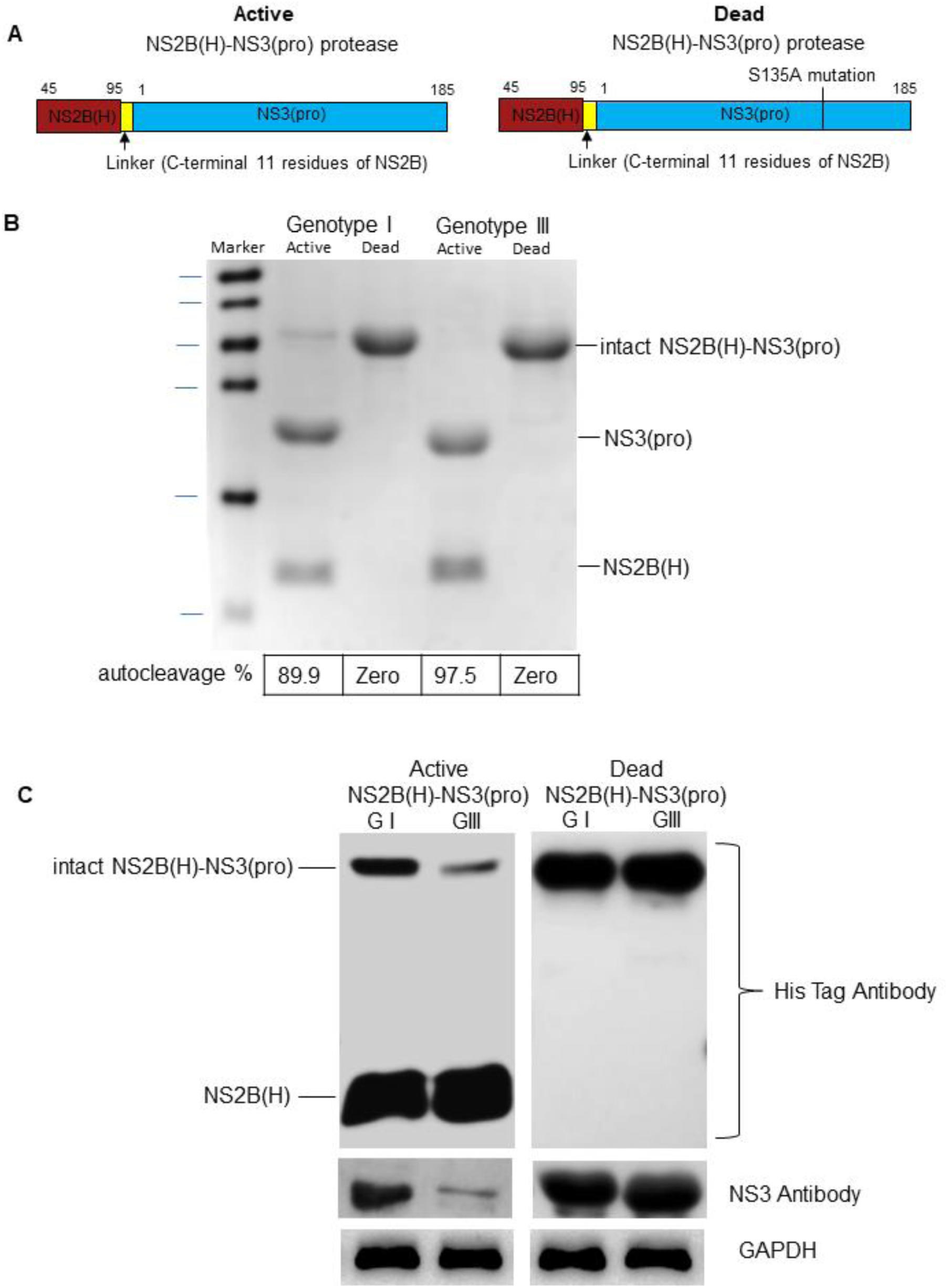
Cloning, expression, and purification of active and dead recombinant NS2B(H)-NS3(pro) proteases. **A)** Schematic representation of recombinant plasmids. **B)** SDS-PAGE for active and dead proteases NS2B(H)-NS3pro of GI and GIII. The Bands were quantified by imageJ software to determine the self cleavage percentage. **C)** Western Blot for active and dead NS2B(H)-NS3 of GI and GIII was performed using His tag, JEV NS3 and GADBH antibodies.

### Analysis of proteolytic processing activities of GI and GIII NS2B(H)-NS3(pro) proteases using fluorogenic model

Enzymatic activity of GI-SH7 and GIII-SH15 recombinant NS2B(H)-NS3(pro) proteases was examined, and compared by fluorescence obtained after hydrolysis of fluorogenic peptide substrates containing the sequence identical to dibasic cleavage sites of NS2A/NS2B and NS2B/NS3 from JEV polyprotein [42, 52].

The assays conducted to compare proteolytic activities of active and inactivated proteases using the fluorogenic peptide containing cleavage site sequence from JEV NS2B-NS3 revealed that all four groups of proteases did not display differences in kinetic data/florescence at zero minutes indicating the non-hydrolysis of the suspectable cleavage site on specific time point. However, after a twenty minute interval a statistically significant difference was seen between GI-SH7 and GIII-SH15 (Fig. 4A). The GI-SH7 and GIII-SH15 also exhibited statistical differences from their respective dead proteases, indicating cleavage of fluorogenic substrate by active proteases of both strains. The kinetic values/fluorescence data kept increasing with passage of time and the values became constant at 480 minutes. Significant differences were observed in all time points between GI-SH7 and GIII-SH15 from forty minutes onwards. The dead proteases fluorescence data remained constant without any increase after zero minutes throughout the experiment (Fig. 4A).

**Figure 4:**
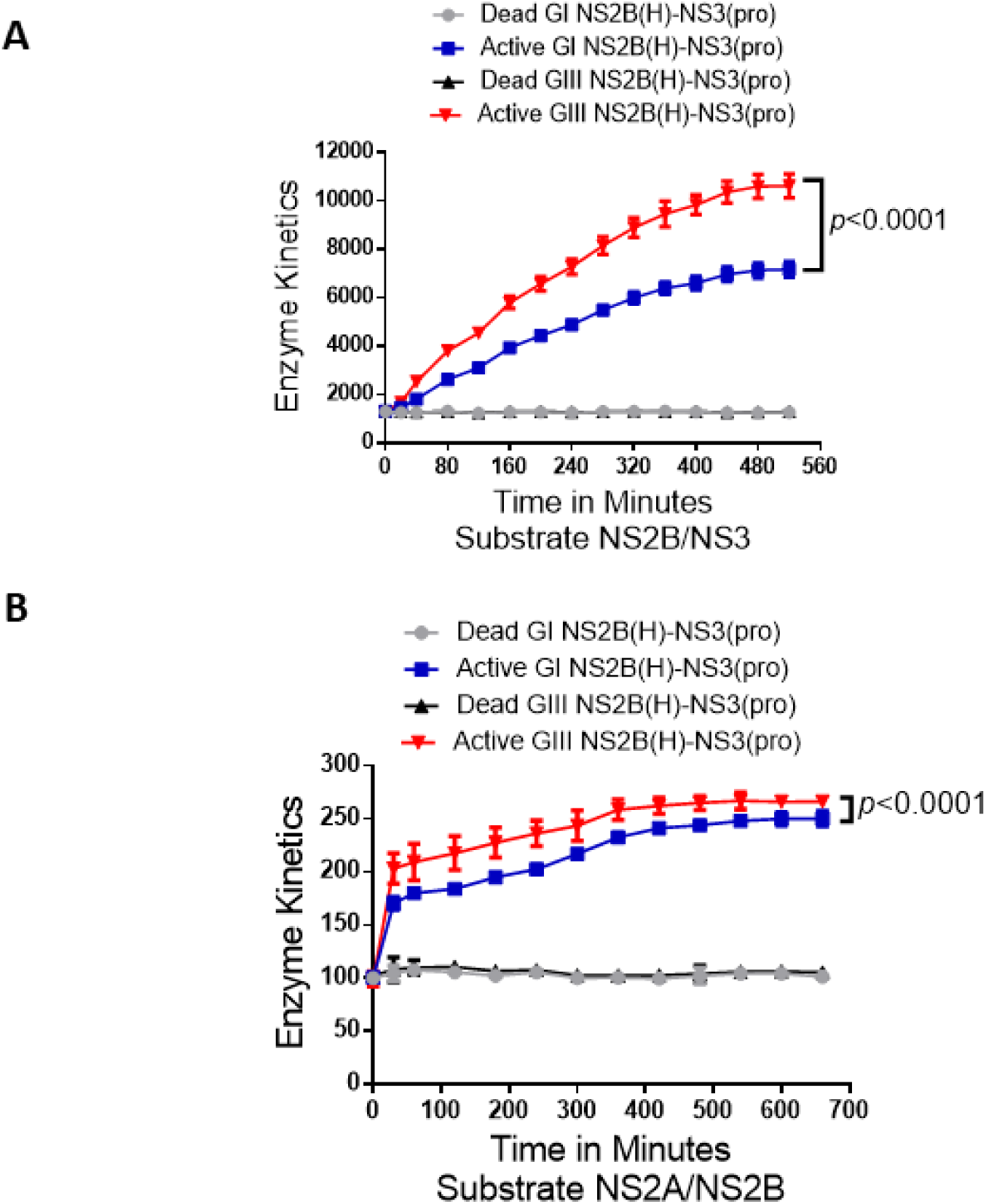
Active and dead JEV NS2B(H)-NS3(pro) catalyzed substrate hydrolysis rates. **A)** The graph shows reaction velocities/enzyme activities difference of active and dead JEV genotype I and III NS2B(H)-NS3(pro) proteases for hydrolysis of Pyr-RTKR-AMC, a substrate site gotten from JEV cleavage site NS2B-NS3. **B)** The graph represents reaction velocities/enzyme activities difference of JEV genotype I and III active and dead NS2B(H)-NS3(pro) proteases for hydrolysis of Dabcyl-PNKKRGWPAT-(Edans)G, a substrate cleavage site gotten from JEV NS2A-NS2B.

The experiments were repeated using synthetic fluorogenic peptides from cleavage sites of JEV NS2A/NS2B. All the four group of proteases did not display any difference in kinetic data at zero minutes. At forty minutes a statically significant difference was seen between GI-SH7 and GIII-SH15 (Fig. 4B). The GI-SH7 and GIII-SH15 proteases also exhibited statistically significant differences from their respective dead/inactive proteases. These kinetic values kept increasing and became constant at 480 minutes. The dead proteases fluorescence data remained constant without any increase throughout experiment indicating non-hydrolysis of fluorogenic substrate by dead proteases (Fig. 4B).

Collectively, these findings revealed that proteases from GIII-SH15 possess high proteolytic processing activities compared to GI-SH7 and the artificial fluorogenic peptide containing cleavage sites from NS2B/NS3 are more efficiently cleaved by JEV proteases compared to site NS2A/NS2B.

### Proteolytic Processing activities of GI and GIII NS2B(H)-NS3(pro) proteases at elevated temperatures

The higher thermal stability at elevated temperatures for GI could be a causative factor related to enhancement of viral replication of GI viruses. Previously, NS2B/NS3 mutations in JEV were found to enhance infectivity of GI over GIII in amplifying host cells at elevated temperatures [31]. To access the influence of viral thermal stability on viral NS2B/NS3 protease proteolytic activities the experiment was repeated using elevated temperatures of 41°C with fluorogenic substrate NS2B/NS3. All viral proteases did not display any hydrolysis of fluorogenic peptide at zero minutes. From twenty minutes onwards a statistically significant difference was seen between GI-SH7 and GIII-SH15 activities (Fig. 4). These kinetic values kept increasing variably (maintaining the significant difference between both genotypes) and become constant at 400 minutes onwards, whilst dead protease fluorescence data remained constant without any increase throughout experiment (Fig. 5). These results were similar to experiments performed at 37 ℃ (Fig. 4) and indicated that elevated temperatures do not influence proteolytic processing activities of viral proteases.

**Figure 5:**
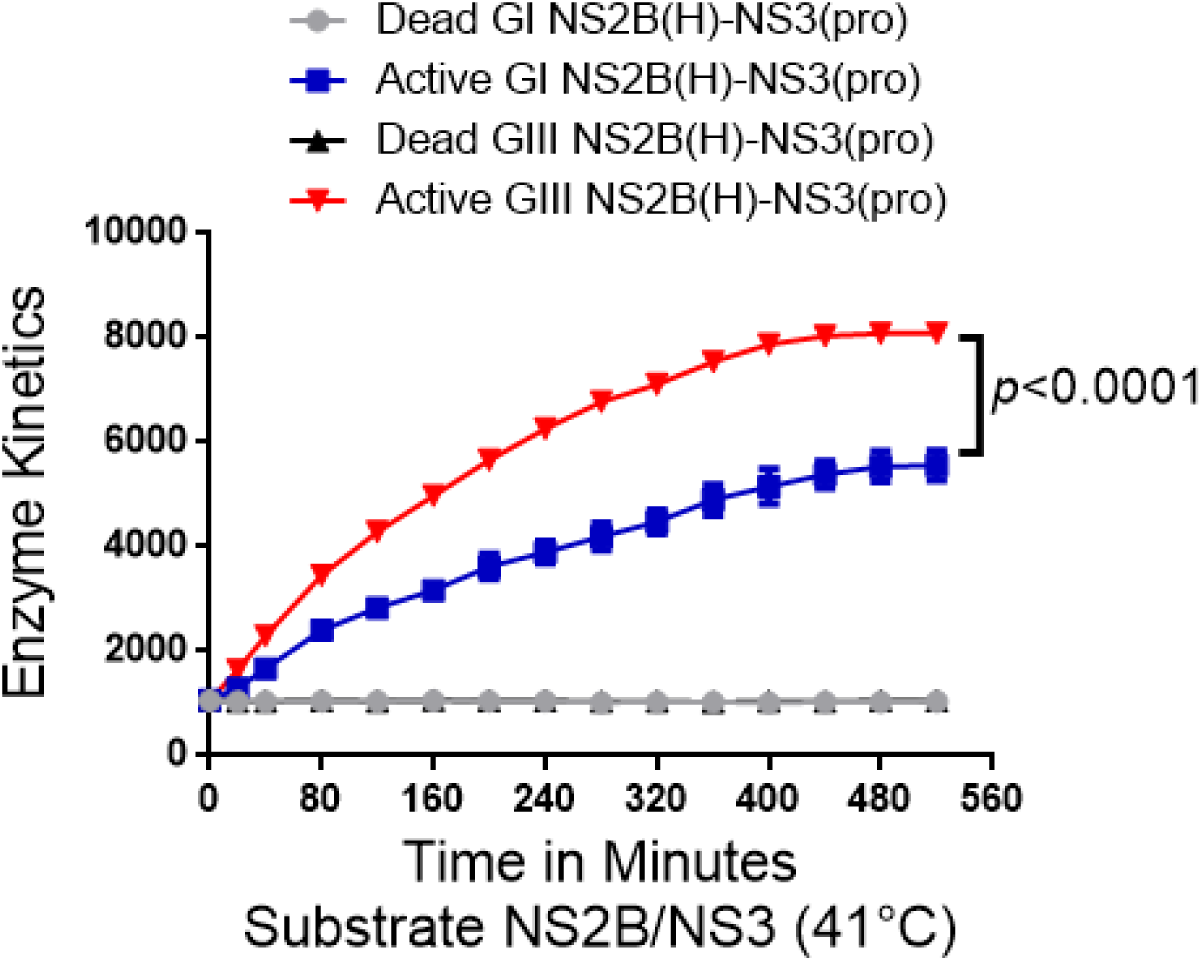
Active and dead JEV NS2B(H)-NS3(pro) catalyzed substrate hydrolysis rates at elevated temperatures. **A)** The graph represents reaction velocities difference of JEV genotype I and III active and dead NS2B(H)-NS3(pro) proteases for hydrolysis of A)Pyr-RTKR-AMC and **B)** Dabcyl-PNKKRGWPAT-(Edans)G, substrate sites gotten from JEV NS2B-NS3 and JEV NS2A/NS2B at elevated temperatures i.e 41C.

### Hydrophilic domain of NS2B determines the difference in NS2B(H)-NS3(pro) protease activities between GI and GIII

As synthetic fluorogenic peptide harbouring cleavage sites from NS2B/NS3 was cleaved more efficiently by JEV proteases so it was selected to determine whether hydrophilic amino acid variations in the protease domain in viral proteases are involved in increased proteolytic processing or hydrolysis of fluorogenic peptides of GIII-SH15 over GI-SH7. Recombinant proteases were generated by exchanging proteins encoding hydrophilic domain from NS2B and protease domain from NS3 of respective strains GI-SH7 and GIII-SH15 strains resulting in GI/NS2B(H)-GIII/NS3(pro) and GIII/NS2B(H)-GI/NS3(pro) (Fig. 6A). Exchanging the hydrophilic domain of GI NS2B to GIII in GI protease (GIII/NS2B(H)-GI/NS3(pro)) significantly increased its proteolytic processing activities as compared with the parental GI NS2B(H)-NS3(pro) protease (Fig. 6B). The activity of GIII/NS2B(H)-GI/NS3(pro) became almost similar to that of GIII NS2B(H)-NS3(pro) with no significant difference among them (Fig. 6C).

**Figure 6:**
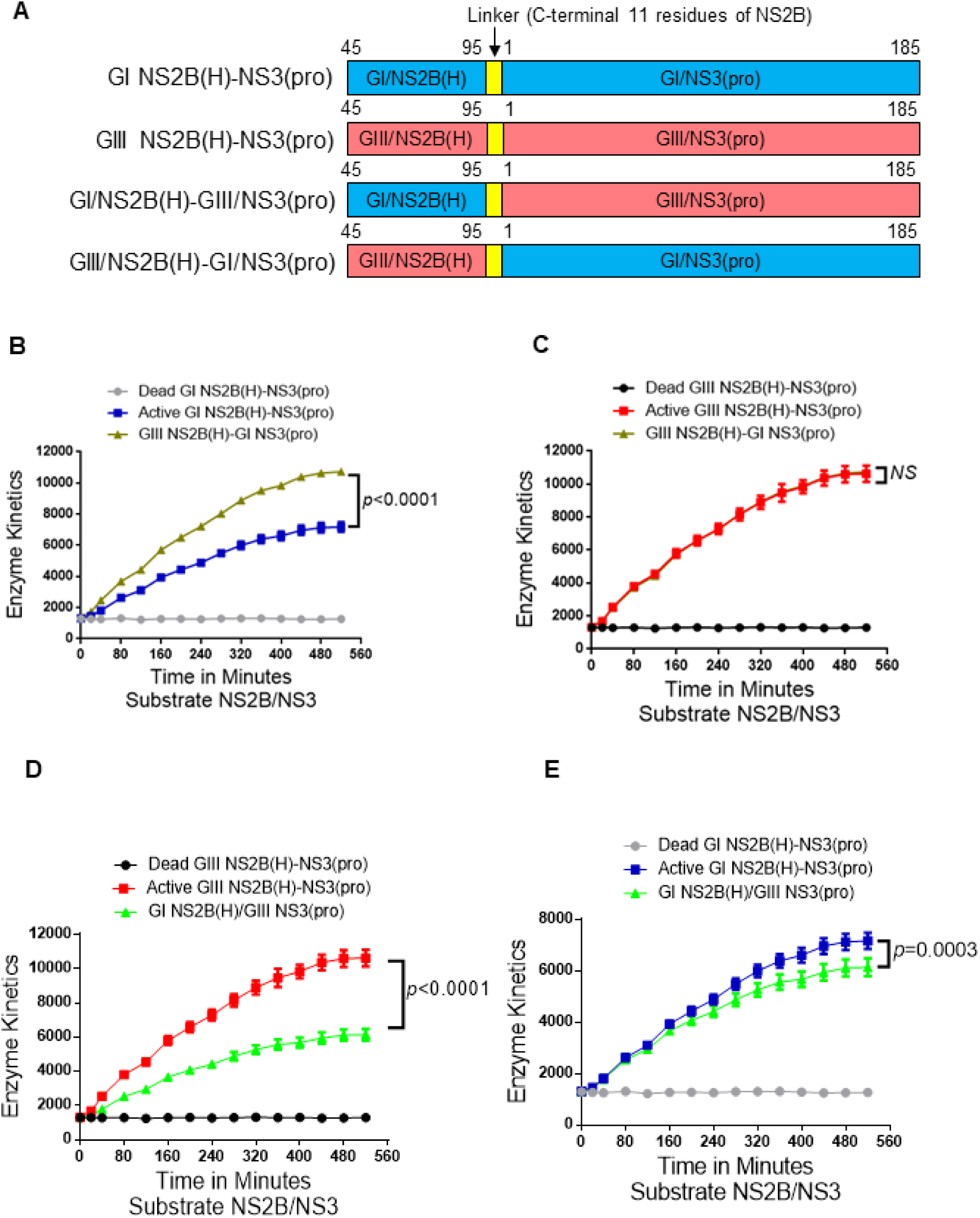
Role of NS2B Hydrophilic domain in differential hydrolysis rates/proteolytic processing activities. **A)** Schematic representation of reversed JEV GI-SH7 i.e GIII NS2B(H)-GI NS3(pro) with exchanged hydrophilic domain from GIII-SH15 and reversed JEV GIII-SH15 i.e GI NS2B(H)/GIII NS3(pro) with exchanged hydrophilic domains from GI-SH7. **(B and C)**The graphs represents and compares reaction velocities/enzyme activities difference of JEV genotype GI-SH7 **(B)** and GIII-SH15 **(C)**with reversed JEV GI-SH7 i.e GIII NS2B(H)-GI NS3(pro) for hydrolysis of Pyr-RTKR-AMC, a substrate site gotten from JEV NS2B-NS3**. (D and E)** The graphs represents and compares reaction velocities/enzyme activities difference of JEV genotype GIII-SH15 **(D)** and GI-SH7 **(E)** with reversed JEV GIII-SH15 i.e GI NS2B(H)-GIII NS3(pro) for hydrolysis of Pyr-RTKR-AMC, a substrate site gotten from JEV NS2B-NS3.

After exchanging the hydrophilic domain of GIII NS2B to GI in GIII proteases (GI/NS2B(H)-GIII/NS3(pro)), a significant decrease in the activity of GI/NS2B(H)-GIII/NS3(pro) protease was observed compared to the parental GIII NS2B(H)-NS3(pro) protease (Fig. 6D). The levels of GI/NS2B(H)-GIII/NS3(pro) protease activity were similar to those of GI NS2B(H)-NS3(pro) protease during 20-120 min, but significantly lower than those of GI NS2B(H)-NS3(pro) protease from 160 min (Fig. 6E). Collectively, these results demonstrate that the hydrophilic domain of NS2B determined the difference in NS2B(H)-NS3(pro) protease activities between GI and GIII, suggesting that the mutations in the hydrophilic domain of NS2B may be responsible for the difference in NS2B(H)-NS3(pro) protease activities between GI and GIII.

### Contribution of NS2B-D55E variation in hydrophilic domain of NS2B to the difference in NS2B(H)-NS3(pro) protease activities between GI and GIII

The hydrophilic domain of NS2B is responsible for differences in proteolytic processing activities between GI and GIII NS2B(H)-NS3(pro) proteases. This specific domain possesses two variations at position 55 (NS2B-D55E) and 65 (NS2B-E65D) in the NS2B protein (Fig. 1C). Data obtained after alignment of fifty represented strains of GI and GIII revealed that both mutations have conservation rate of 90%-100% [49]. To determine the contribution of NS2B-D55E to differential proteolytic processing activities between GI and GIII NS2B(H)-NS3(pro) proteases, we replaced aspartic acid (D) at position 55 of GI NS2B(H)-NS3(pro) protease with glutamic acid (E) to generate a mutant protease of GI NS2B(H)/D55E-NS3(pro) and substituted aspartic acid (D) for glutamic acid (E) at position 55 of GIII NS2B(H)-NS3(pro) protease to generate a mutant protease of GIII NS2B(H)/E55D-NS3(pro) (Fig. 7A). Substitution of NS2B-D55E variation resulted in a conformational change of NS2B(H), as compared with its respective parent (Fig. 7B). The mutants of GI NS2B(H)/D55E-NS3(pro) and GIII NS2B(H)/E55D-NS3(pro) were prepared, and their proteolytic processing activities compared with their parental GI NS2B(H)-NS3(pro) and GIII NS2B(H)-NS3(pro) proteases. Exchanging the amino acid residue at 55 in GI NS2B(H) from aspartic acid to glutamic acid significantly increased the proteolytic processing activities of GI NS2B(H)/D55E-NS3(pro), as compared to its parental GI NS2B(H)-NS3(pro) protease (Fig. 7C). However, the levels of GI NS2B(H)/D55E-NS3(pro) proteas activities remained lesser than those of GIII NS2B(H)-NS3(pro) protease (Fig. 7D). These results indicate that substitution of NS2B-D55E contributed to the increased proteolytic processing of GI NS2B(H)/D55E-NS3(pro) over its parental GI NS2B(H)-NS3(pro). When the amino acid at position 55 in GIII NS2B(H) was exchanged from glutamic acid to aspartic acid, the levels of GIII NS2B(H)/E55D-NS3(pro) protease activities significantly decreased, as compared to its parental GIII NS2B(H)-NS3(pro) protease (Fig. 7E), but remained higher than those of GI NS2B(H)-NS3(pro) protease (Fig. 7F). Collectively, these results demonstrated that the NS2B-D55E variation in hydrophilic domain of NS2B contributed to the difference in NS2B(H)-NS3(pro) protease activities between GI and GIII.

**Figure 7:**
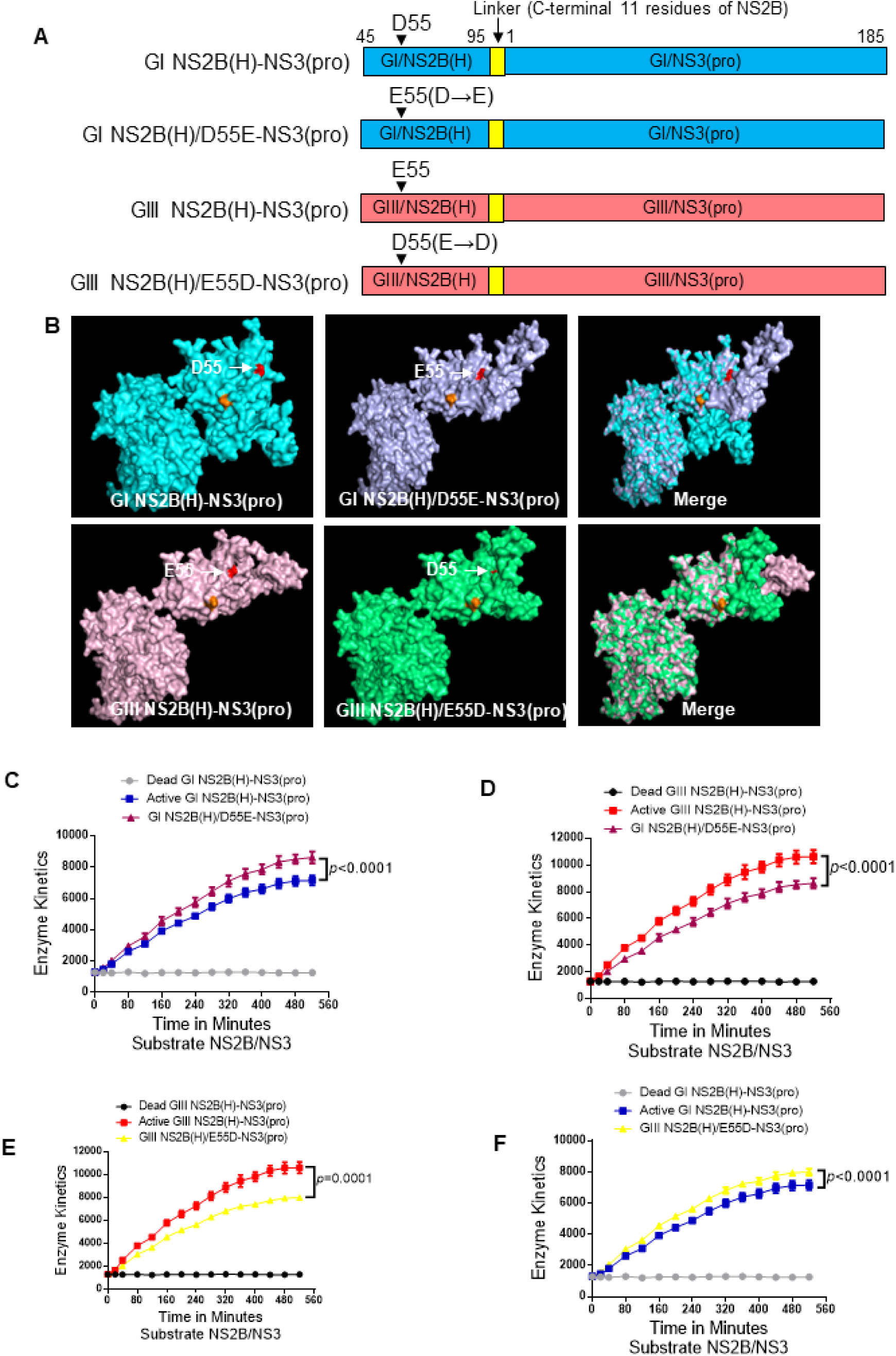
Contribution of variation 55 in hydrophilic domain of NS2B to proteolytic activities of NS2B(H)-NS3(pro). **A).** Schematic diagram of reversed JEV GI-SH7 i.e GI NS2B(H)/D55E-NS3(pro) and reversed JEV GIII-SH15 i.e GIII NS2B(H)/E55D-NS3(pro) with exchanged NS2B-55 variations from their respective parent NS2B(H)-NS3(pro) proteases. **B)** The graph represents and compares 3D structural conformations of reversed JEV GI-SH7 i.e GI-SH7 i.e GI NS2B(H)/D55E-NS3(pro) and reversed JEV GIII-SH15 i.e GIII NS2B(H)/E55D-NS3(pro) with their respective parent GI-SH7 and GIII-SH15 NS2B(H)-NS3(pro) proteases. Variation NS2B(H)55 is highlighted in red. **C and D)** The graphs represents and compares reaction velocities/enzyme activities difference of JEV genotype GI-SH7 **(C)** and GIII-SH15 NS2B(H)-NS3(pro) proteases **(D)** with reversed JEV GI-SH7 i.e GI NS2B(H)/D55E-NS3(pro) for hydrolysis of Pyr-RTKR-AMC, a substrate site gotten from JEV NS2B-NS3. **E and F)** The graphs represents and compares reaction velocities/enzyme activities difference of JEV genotype GI-SH7 **(F)** and GIII-SH15 **(E)** NS2B(H)-NS3(pro) proteases with reversed JEV GIII-SH15 i.e GIII NS2B(H)/E55D-NS3(pro) for hydrolysis of Pyr-RTKR-AMC, a substrate site gotten from JEV NS2B-NS3

### Contribution of NS2B-E65D variation in hydrophilic domain of NS2B to the difference in NS2B(H)-NS3(pro) protease activities between GI and GIII

To determine the contribution of NS2B-E65D variation in hydrophilic domain of NS2B to the difference in NS2B(H)-NS3(pro) protease activities between GI and GIII, we replaced glutamic acid (E) at position 65 of GI NS2B(H)-NS3(pro) protease with aspartic acid (D) to generate a mutant protease of GI NS2B(H)/E65D-NS3(pro) and substituted glutamic acid (E) for aspartic acid (D) at position 65 of GIII NS2B(H)-NS3(pro) protease to generate a mutant protease of GIII NS2B(H)/D65E-NS3(pro) (Fig. 8A). Substitution of NS2B-E65D variation resulted in a conformational change of NS2B(H), as compared with its respective parent (Fig. 8B). The mutants of GI NS2B(H)/E65D-NS3(pro) and GIII NS2B(H)/D65E-NS3(pro) were prepared, and their proteolytic processing activities were compared with their parental GI NS2B(H)-NS3(pro) and GIII NS2B(H)-NS3(pro) proteases. Exchanging the amino acid residue at 65 in GI NS2B(H) from glutamic acid to aspartic acid significantly increased the proteolytic processing activities of GI NS2B(H)/E65D-NS3(pro), as compared to its parental GI NS2B(H)-NS3(pro) protease (Fig. 8C). However, the levels of GI NS2B(H)/E65D-NS3(pro) proteas activities remained lesser than those of GIII NS2B(H)-NS3(pro) protease (Fig. 8D). These results indicated that co-substitution of NS2B-E65D contributed to the increased proteolytic processing of GI NS2B(H)/E65D-NS3(pro) over its parental GI NS2B(H)-NS3(pro). When the amino acid residue at position 65 in GIII NS2B(H) was exchanged from aspartic acid to glutamic acid, the levels of GIII NS2B(H)/D65E-NS3(pro) protease activities significantly decreased, as compared to its parental GIII NS2B(H)-NS3(pro) protease (Fig. 8E), but remained higher than those of GI NS2B(H)-NS3(pro) protease (Fig. 8F). Collectively, these results demonstrated that the NS2B-E65D variation in hydrophilic domain of NS2B contributed to the difference in NS2B(H)-NS3(pro) protease activities between GI and GIII.

**Figure 8:**
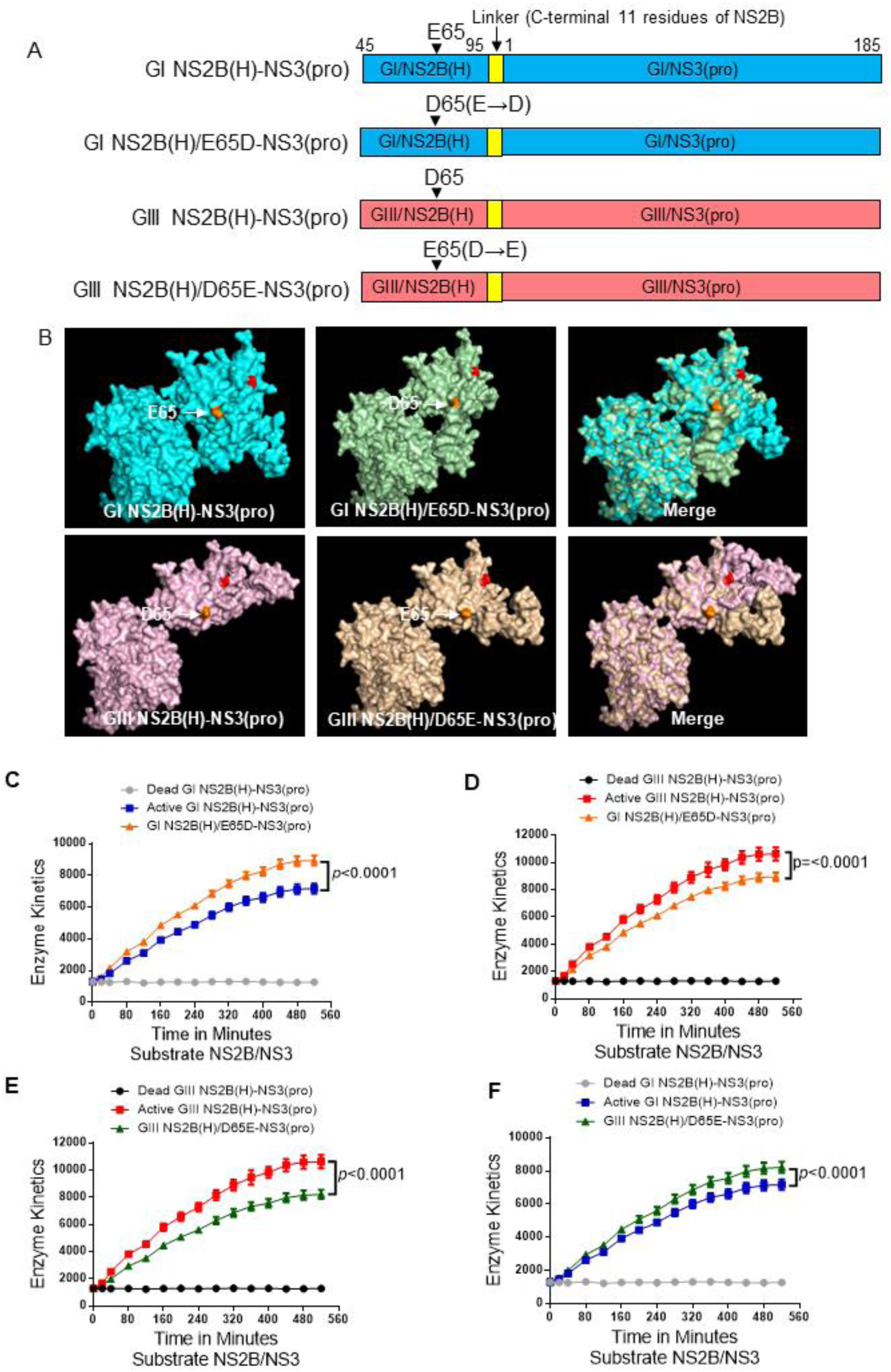
Contribution of variation 65 in hydrophilic domain of NS2B to proteolytic activities of NS2B(H)-NS3(pro). **A).** Schematic diagram of reversed JEV GI-SH7 i.e GI NS2B(H)/ E65D-NS3(pro) and reversed JEV GIII-SH15 i.e GIII NS2B(H)/D65E-NS3(pro) with exchanged NS2B-65 variations from their respective parent NS2B(H)-NS3(pro) proteases. **B)** The graph represents and compares 3D structural conformations of reversed JEV GI-SH7 i.e GI NS2B(H)/ E65D-NS3(pro) and reversed JEV GIII-SH15 i.e GIII NS2B(H)/D65E-NS3(pro) with their respective parent GI-SH7 and GIII-SH15 NS2B(H)-NS3(pro) proteases. Variation NS2B(H)65 is highlighted in yellow. **C and D)** The graphs represents and compares reaction velocities/enzyme activities difference of JEV genotype GI-SH7 **(C)** and GIII-SH15 NS2B(H)-NS3(pro) proteases **(D)** with reversed JEV GI-SH7 i.e GI NS2B(H)/ E65D-NS3(pro) for hydrolysis of Pyr-RTKR-AMC, a substrate site gotten from JEV NS2B-NS3. **E and F)** The graphs represents and compares reaction velocities/enzyme activities difference of JEV genotype GI-SH7 **(F)** and GIII-SH15 **(E)** NS2B(H)-NS3(pro) proteases with reversed JEV GIII-SH15 i.e GIII NS2B(H)/D65E-NS3(pro) for hydrolysis of Pyr-RTKR-AMC, a substrate site gotten from JEV NS2B-NS3.

### Co-contribution of NS2B-D55E and NS2B-E65D variations in hydrophilic domain of NS2B to the difference in NS2B(H)-NS3(pro) protease activities between GI and GIII

Variation in the hydrophilic domain of both NS2B-D55E and NS2B-E65D contributed individually to the difference in NS2B(H)-NS3(pro) protease activities between GI and GIII. The amino acid residues at position 55 and 65 of NS2B were simultaneously exchanged between GI and GIII NS2B(H)-NS3(pro) proteases to generate the co-substitution mutants of GI NS2B(H)/D55E/E65D-NS3(pro) and GIII NS2B(H)/E55D/D65E-NS3(pro) (Fig. 9A). Co-substitution of NS2B-D55E and NS2B-E65D resulted in a conformational change of NS2B(H), as compared with its respective parent (Fig. 9B). Analysis of the proteolytic processing activity indicated that co-substitution of NS2B-D55E and NS2B-E65D significantly increased the levels of proteolytic processing activity of GI NS2B(H)/D55E/E65D-NS3(pro), as compared to those of its parental GI NS2B(H)-NS3(pro) (Fig. 9C), the levels increased by co-substitution were similar to those of GIII NS2B(H)-NS3(pro) (Fig. 9D). Together these results indicated that co-substitution of NS2B-D55E and NS2B-E65D contributed collectively to the increased proteolytic processing of GI NS2B(H)/D55E/E65D-NS3(pro) over its parental GI NS2B(H)-NS3(pro). On another hand, the co-substitution of NS2B-E55D and NS2B-D65E altered the proteolytic processing activity of GIII NS2B(H)-NS3(pro). The levels of proteolytic processing activity of GIII NS2B(H)/E55D/D65E-NS3(pro) were significantly lower than those of its parental GIII NS2B(H)-NS3(pro) (Fig. 9E) as well as relatively lower than those of GI NS2B(H)-NS3(pro) (Fig. 9F). Overall, these data suggested that NS2B-D55E and NS2B-E65D variations in hydrophilic domain of NS2B co-contributed to the difference in NS2B(H)-NS3(pro) protease activities between GI and GIII.

**Figure 9:**
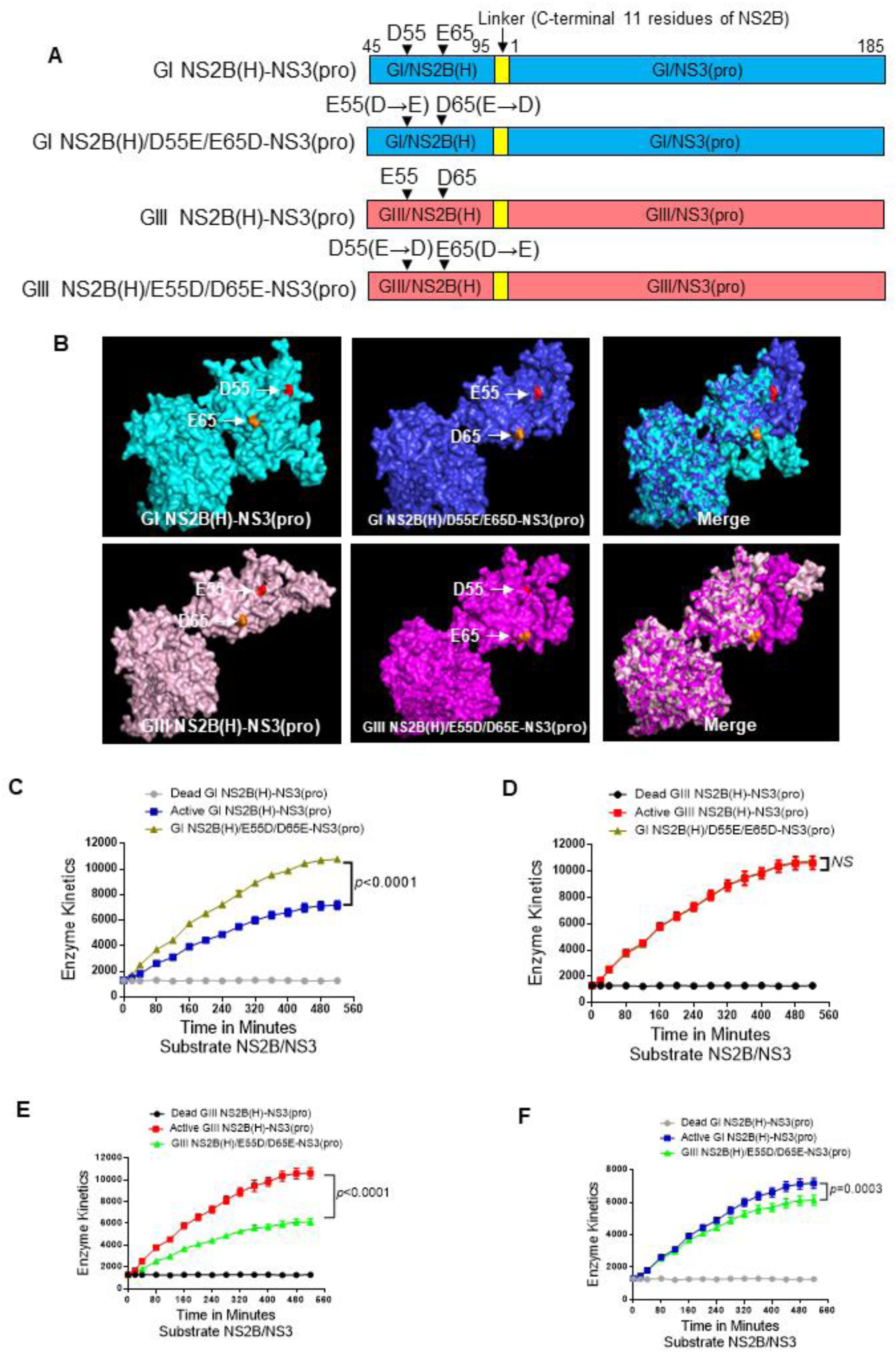
Co-contribution of NS2B-D55E and NS2B-E65D variations in hydrophilic domain of NS2B to proteolytic activities of NS2B(H)-NS3(pro). **A).** Schematic diagram of reversed JEV GI-SH7 i.e GI NS2B(H)/D55E/E65D-NS3(pro) and reversed JEV GIII-SH15 i.e GIII NS2B(H)/E55D/D65E-NS3(pro) with exchanged NS2B-D55E and NS2B-E65D variations from their respective parent NS2B(H)-NS3(pro) proteases. **B)** The graph represents and compares 3D structural conformations of reversed JEV GI-SH7 i.e GI NS2B(H)/D55E/E65D-NS3(pro) and reversed JEV GIII-SH15 i.e GIII NS2B(H)/E55D/D65E-NS3(pro) with their respective parent GI-SH7 and GIII-SH15 NS2B(H)-NS3(pro) proteases. Variation NS2B(H)55 and NS2B(H) 65 are highlighted in red and yellow respectively. **C and D)** The graphs represents and compares reaction velocities/enzyme activities difference of JEV genotype GI-SH7 **(C)** and GIII-SH15 NS2B(H)-NS3(pro) proteases **(D)** with reversed JEV GI-SH7 i.e GI NS2B(H)/D55E/E65D-NS3(pro) for hydrolysis of Pyr-RTKR-AMC, a substrate site gotten from JEV NS2B-NS3 **E and F)** The graphs represents and compares reaction velocities/enzyme activities difference of JEV genotype GI-SH7 **(F)** and GIII-SH15 **(E)** NS2B(H)-NS3(pro) proteases with reversed JEV GIII-SH15 i.e GIII NS2B(H)/E55D/D65E-NS3(pro) for hydrolysis of Pyr-RTKR-AMC, a substrate site gotten from JEV NS2B-NS3.

## Discussion

Emerging JEV GI virus has gradually displaced JEV GIII virus as a dominant virus genotype isolated from stillborn piglets, *Culex tritaeniorhynchus* and human cases since 1990s. The mechanism behind this genotype replacement still remains unclear. In this study we have identified the contribution of JEV NS2B(H)-NS3(pro) proteases determinants which are corelated with enhanced proteolytic processing activities of GIII proteases using fluorogenic peptide substrate model.

JEV polyprotein is cleaved to generate functional proteins by complex combination of host and viral proteases. The cleavage is predicted to occur at junctions between C/prM, prM/E, E/NS1, NS1/NS2A, NS2B/NS3, NS3/NS4A, NS4A/NS4B, NS4B/NS5 and sites of internal C, NS4A and NS3 [1, 2]. Studies have demonstrated that JEV protease activity mainly depends on association between NS3 and its co-factor NS2B, and that this two component NS2B-NS3 proteases expressed in *E.coli* are folded correctly with effective proteolytic activity. [28, 42].

We found that GI-SH7 exhibits different 3D protein conformations (Figure 7B) and eight critical conserved substitutions (Figure 1C) in NS2B(H)-NS3(pro) protease region than GIII-SH15. Data obtained after alignment of fifty represented strains of GI and GIII revealed that two mutations in NS2B hydrophilic domain have a conservation rate of 90%-100%, Whereas four mutations in NS3 proteases are conserved with rate of 90-100 % and two with conservation rate of 50-89% [49]. Some previous observations have suggested that differences in the conformational space between NS2B-NS3 proteases of different flaviviruses might lead to the differences in substrate recognition and affinity, [52–55] [56, 57]. To analyze and compare the cleavage pattern of GI-SH7 with GIII-SH15 NS2B/NS3 proteases, the active and inactivated GI-SH7 and GIII-Sh15 [44] NS2B(H)-NS3(pro) proteases were structured and engineered into PET-Duet 1 vector for expression in *E.coli* and cleavage pattern was assessed by western blot using respective antibodies (Figure 2). Our results demonstrated that GI-SH7 proteases were able to cleave the sites at internal C, NS2A/NS2B, NS2B/NS3, and NS3/NS4A junctions and exhibited the same cleavage pattern as previously found for SH-GIII proteases [51], suggesting that viral protease substitution and differential protein conformation do not interfere with selection of cleavage sites for proteolytic processing and possibly play no role in genotype displacement in this specific side. The identical cleavage patterns of both genotypes comprising critical mutations were supported by previous studies demonstrating that substrate recognizing sequence is highly conserved among all flaviviruses and contains two basic residues in P2 and P1 followed by a small unbranched amino acid in P1’ [58, 59].

To analyze and compare the proteolytic processing activities of GI-SH7 and GIII-SH15 NS2B/NS3 proteases, the active and inactivated NS2B(H)-NS3(pro) proteases were structured and engineered into P-TrcHisA vector for expression in *E.coli* (Fig. 3A). SDS-PAGE of purified proteins revealed presence of three bands at different sizes indicating autocleavage and only the intact NS2B(H)-NS3(pro) proteases band for inactivated/dead NS2B/NS3 proteases confirming its inactivity. Quantification of blots revealed that SH7 was self-cleaved with a percentage 89.9% of and SH15 with a percentage of 97.5%. Obvious differences in intact anti-His and anti-NS3 expression was also seen between both strains (Fig. 3C). These results suggested that GIII-SH15 shows high autocleavage rate of enzymatically active NS2B(H)-NS3(pro) proteases as compared to GI-SH7. The autocleavage abilities of various flaviviruses proteases have also been reported in previous studies [42, 43, 56].

Enzymatic activity of GI-SH7 and GIII-SH15 recombinant NS2B(H)-NS3(pro) proteases was examined and compared by fluorescence obtained after hydrolysis of fluorogenic peptide substrates containing the sequence identical to dibasic cleavage sites of NS2A/NS2B and NS2B/NS3 from JEV polyprotein. Data revealed that proteases from GIII-SH15 possess high proteolytic processing activities when compared to GI-SH7 and the artificial fluorogenic peptide containing cleavage site from NS2B/NS3(Fig 4A) are more efficiently cleaved by JEV proteases as compared to site NS2A/NS2B (Fig 4B). The higher thermal stability at elevated temperatures for GI could be a causative factor related to enhancement of viral replication of GI viruses. Previously, NS2B/NS3 mutations in Japanese encephalitis virus were found to enhance infectivity of GI over GIII in amplifying host cells at elevated temperatures [31]. To access the influence of viral thermal stability on viral NS2B/NS3 proteases proteolytic activities the experiment was repeated using elevated temperatures of 41°C with fluorogenic substrate NS2B/NS3. The trends were identical to experiments performed at 37°C (Fig. 5) and indicated that elevated temperatures don’t influence proteolytic processing activities of viral proteases. And GIII-SH15 possess high proteolytic activities than GI-SH7 at normal and elevated temperatures.

In order to determine whether amino acid variations of hydrophilic or protease domain in viral proteases are responsible and involved in increased proteolytic processing of GIII-SH15. The recombinant proteases were generated by exchanging proteins encoding hydrophilic domain from NS2B and protease domain from NS3 of respective strains GI-SH7 and GIII-SH15 strains resulting in GI/NS2B(H)-GIII/NS3(pro) and GIII/NS2B(H)-GI/NS3(pro) (Fig. 6A). Exchanging the hydrophilic domain of GI NS2B to GIII in GI protease (GIII/NS2B(H)-GI/NS3(pro)) significantly increased it proteolytic processing activities, as compared with its parental GI protease (Fig. 6B). The activity of recombinant proteases GIII/NS2B(H)-GI/NS3(pro) became almost similar to that of GIII proteases with non-significant difference among them (Fig. 6C). On the other hand, after exchanging the hydrophilic domain of GIII NS2B to GI in GIII proteases (GI/NS2B(H)-GIII/NS3(pro)), a significant decrease in its activity was observed, compared to its parental GIII proteases (Fig. 6D). However, the levels of GI/NS2B(H)-GIII/NS3(pro) protease activity become significantly lower than those of GI protease (Fig. 6E). Collectively, these results demonstrated that the hydrophilic domain of NS2B determined the difference in NS2B(H)-NS3(pro) protease activities between GI and GIII, suggesting that the mutations in the hydrophilic domain of NS2B may be responsible for the difference in NS2B(H)-NS3(pro) protease activities between GI and GIII. This specific domain possesses two conserved variations at position 55 (NS2B-D55E) and 65 (NS2B-E65D) in NS2B protein (Fig. 1C) [49]. To determine the contribution of NS2B-D55E to differential proteolytic processing activities between GI and GIII, we replaced aspartic acid (D) at position 55 of GI proteases protease with glutamic acid (E) to generate a mutant protease of GI NS2B(H)/D55E-NS3(pro) and substituted aspartic acid (D) for glutamic acid (E) at position 55 of GIII proteases to generate a mutant protease of GIII NS2B(H)/E55D-NS3(pro) (Fig. 7A). NS2B-D55E and NS2B-E55D variation resulted in a conformational change of NS2B(H), as compared with their respective parent strains (Fig. 7B). Exchanging the amino acid residue at 55 in GI NS2B(H) significantly increased the proteolytic processing activities of GI NS2B(H)/D55E-NS3(pro), as compared to its parental GI protease (Fig. 7C). However, the levels of GI NS2B(H)/D55E-NS3(pro) proteases activities remained lesser than those of GIII (Fig. 7D) indicating contribution of NS2B-D55E to increase proteolytic processing. When the amino acid at position 55 in GIII NS2B(H) was exchanged from glutamic acid to aspartic acid, the levels of GIII NS2B(H)/E55D-NS3(pro) protease activities significantly decreased, as compared to its parental GIII protease (Fig. 7E) but remained higher than those of GI protease (Fig. 7F). Collectively, these results demonstrated that the NS2B-D55E variation in hydrophilic domain of NS2B contributes to the difference in NS2B(H)-NS3(pro) protease activities between GI and GIII. To determine the contribution of NS2B-E65D variation to differential proteolytic processing activities between GI and GIII we replaced glutamic acid (E) at position 65 of GI protease with aspartic acid (D) to generate a mutant protease of GI NS2B(H)/E65D-NS3(pro) and substituted glutamic acid (E) for aspartic acid (D) at position 65 of GIII proteases to generate a mutant protease of GIII NS2B(H)/D65E-NS3(pro) (Fig. 8A). Substitution of NS2B-E65D NS2B-D65E variations resulted in a conformational change of NS2B(H), as compared with their respective parent strains (Fig. 8B). Exchanging the amino acid residue at 65 in GI NS2B(H) significantly increased the proteolytic processing activities of GI NS2B(H)/E65D-NS3(pro), as compared to its parental GI protease (Fig. 8C). However, the levels of GI NS2B(H)/E65D-NS3(pro) proteas activities remained lesser than those of GIII protease (Fig. 8D) indicating contribution of NS2B-D55E to increase proteolytic processing. When the amino acid residue at position 65 in GIII NS2B(H) was exchanged, the levels of GIII NS2B(H)/D65E-NS3(pro) protease activities significantly decreased, as compared to its parental GIII proteases (Fig. 8E), but remained higher than those of GI proteases (Fig. 8F).These results demonstrated that the NS2B-E65D variation in hydrophilic domain of NS2B contributed to the difference in NS2B(H)-NS3(pro) protease activities between GI and GIII. Collectively, these finding indicated that both conserved mutations at position 55 and 65 in hydrophilic domain of NS2B contributes individually and together in increased proteolytic processing activities of GIII proteases over GI in fluorogenic peptide model. These results are braced by previous finding which demonstrates that NS3 protease activity critically depend or controlled by a small NS2B cofactor protein [60, 61] and that the presence of small activating protein or Co-factor is prerequisite for optimal catalytical activity of flavivirus proteases with natural polyprotein substrates[62, 63]. These two mutations may possibly involve in replication advantage of GI over GIII and can provide new insights into molecular basis of JEV genotype shift.

## Conclusion

In conclusion, this study uncovers critical insights into the phenomena of Japanese Encephalitis Virus genotype shifts by identifying and analyzing the specific mutations within the viral NS2B/NS3 proteases. Through *in vitro* analysis, we have revealed genetic determinants in Hydrophilic domain of NS2B/NS3 proteases which may play a pivotal role in genotype shifts and viral replication. Importantly, our research revealed significant differences in protease activities between JEV GI and GIII linked to these mutations, underscoring their functional importance. However, it is crucial to acknowledge that while our study provides essential groundwork, further investigations of observed variations in NS2B/NS3 protease activities using *in vivo* models are important. Exploring these mutations in different animal models or mosquito vectors will provide a deeper understanding of their biological implication and contribute vital data for understanding viral replication, pathogenesis, and development of targeted therapies. This research marks significant enhancement in understanding of JEV genotype shift, emphasizing the need for continued research attempts, enhancing our capacity to combat the varied and dynamic nature of Japanese encephalitis virus and related flaviviruses.

## Funding

The study was supported by the National Natural Science Foundation of China (No. 32273096) awarded to Z.M. The Shanghai Municipal Science and Technology Major Project (No. ZD2021CY001) awarded to Z.M. The Project of Shanghai Science and Technology Commission (No. 22N41900400) awarded to Z.M. The National Key Research and Development Program of China (No. 2022 YFD 1800100) awarded to Y.Q and The project of Cooperation on Animal Biosecurity Prevention and Control in Lancang and Mekong Countries (No. 125161035) awarded to J.W. JLR was supported by NIH grant R01AI12820.

## Data Availability

Relevant data is included within the manuscript.

## Conflict of Interest

The authors declare no conflict of interest.

## Notes

### Competing Interest Statement

The authors have declared no competing interest.

